# Callose deposition at plasmodesmata suggests a role in viral movement restriction in transgenic tobacco plants during recovery from *Tobacco etch virus* infection

**DOI:** 10.1101/2022.11.08.515744

**Authors:** Pablo Vargas-Mejía, Alejandro Olguín-Lamas, Selene L. Fernandez-Valverde, Carmen M. Flores-García, Gertrud Lund, Jean-Philippe Vielle-Calzada, Laura Silva-Rosales

**Author notes:** Correspondence Laura Silva.

## Abstract

Viruses are amongst the most prevalent pathogens that threaten plants. Plants have evolved a sequence-specific defense mechanism against viruses to ensure survival, known as RNA silencing, which includes transcriptional and post-transcriptional gene silencing. After a viral infection, some plants undergo recovery and become further resistant to viral infection. To identify additional mechanisms underlying disease recovery besides the known RNA silencing, we analyzed transgenic tobacco plants expressing a transcript derived from the Nuclear Inclusion “a” protein (NIa) cistron of the *tobacco etch virus* (TEV), which had recovered from infection three weeks following viral inoculation. Using *in situ* hybridizations and qRT-PCR, we detected the viral RNA and the transgene-derived transcript in stem sections adjacent to the recovered leaves. To further characterize the silenced and non-silenced conditions, we undertook tissue-specific RNA-Seq and small RNA-Seq analyses in leaf and stem. We found more differentially expressed genes (DEGs) in the recovered leaf, primarily related to defense, silencing, and hormone signaling responses. Finally, we observed differences in plasmodesmata callose deposition and callose-related genes. Overall, our findings suggest that cell-to-cell viral restriction movement also participates in the recovery of TEV infection in transgenic tobacco plants, besides the key function of RNA silencing.

## 1. Introduction

Viruses, together with fungi and bacteria, are among the most prevalent plant pathogens. Plants have developed specialized strategies to ensure survival in response to constant biotic and abiotic threats. Such strategies include innate and adaptive immune responses involving systemic acquired resistance (SAR) components and RNA interference (RNAi) machinery, respectively [1,2]. One of the possible outcomes of viral infection in plants is symptom recovery, characterized by the emergence of asymptomatic leaves at the apex after systemic symptomatic infection occurrence. Such recovered leaves are virus-free or present a reduction in virus titers and remain resistant to new inoculation of the same or highly related viruses. Recovery from viral disease was first documented in 1928 [1]; later, in 1993, it was linked to the RNA silencing machinery [2–4]. Recently, RNA silencing was shown to be the primary mechanism of viral recovering leaves [3]. Recovery occurs naturally in wild-type plants, especially those infected with positive-sense single-stranded (ss+) RNA viruses and DNA viruses like geminivirus, as well as in those transgenic plants expressing coding or non-coding sequences of the virus genome. Multiple studies have shown that infectious virus is not eliminated during the natural symptom recovery of wild-type plants induced by RNA viruses [5–7].

RNA silencing is a conserved sequence-specific defense mechanism against viruses that has been coopted as an endogenous gene regulatory mechanism. Silencing includes transcriptional (TGS) or post-transcriptional gene silencing (PTGS) [8,9]. PTGS results in the production of small RNAs with the participation of RNA-dependent RNA polymerases (RDRs) producing dsRNA, which are recognized and cleaved by Dicer-like 2 and 4 (DCL2, DCL4) [10,11]. These small dsRNAs can also arise from secondary RNA structures formed during viral genome replication. The resulting 21 to 24 nt cleavage small interfering RNAs (siRNA) are then loaded onto Argonaute proteins AGO1 and AGO2, which form part of the RNA-induced silencing complex (RISC). These AGO-associated siRNAs identify and enable the cleavage of targets through sequence complementarity [12]. RDRs can further amplify the RNA-silencing signal through the vascular system by synthesizing new dsRNA from aberrant RISC-cleaved RNAs recruited by SGS3 [13,14]. Viruses have evolved proteins that counteract silencing by siRNAs, known as VSRs (Viral Silencing Repressors), which sequester virus-derived small RNAs and inhibit their incorporation into the RISC complex. They can also interfere with the expression of the plant silencing machinery through the inactivation or destabilization of RNAi and proteins or by promoting the expression of host endogenous suppressors of RNA silencing (ESRs) such as Nbrgs-CaM (calmodulin from *Nicotiana benthamiana* repressor of gene silencing). The RNAs of viruses encoding weak VSRs, or transgenic plants producing low levels of transgene-derived transcripts, are complemented with the virus-derived transcripts to surpass the RNA threshold level necessary to trigger recovery [4,6,15].

Viral infection recovery mediated by RNA silencing has been primarily studied in leaves, as this is the site where recovery is visually observed. Although much is known about the long-distance movement of silencing signals across the stem, the role of this organ in phenomena such as recovery remains poorly understood [16]. Here, we investigated the expression levels of RNA silencing genes in the stem and leaf tissues during the final recovery stage after viral infection. Additionally, to gain insights into the mechanisms underlying systemic recovery in transgenic plants following viral infection, we looked for other genes that might be differentially expressed between these two organs to identify additional mechanisms underlying disease recovery besides the known RNA silencing observed in the leaves.

## 2. Materials and methods

### 2.1. Plant transformation

The viral NIa cistron sequence was initially cloned into the BamHI sites of plasmids pTL37/8595 containing both T7 and SP6 promoters [17]. Haploid leaf tissue of *Nicotiana tabacum* cultivars B49 and K326 plants was transformed via *Agrobacterium tumefaciens* [18]. A broad host range plasmid pPEV6 containing T-DNA borders, NIa gene, and the neomycin phosphotransferase II gene (that confers kanamycin resistance) selectable marker were mobilized from *E. coli* into *A. tumefaciens,* as described previously by Smith et al. (1994), and used to transform tobacco tissue. Transformed tissue was selected in the presence of 100 µg/mL of kanamycin sulfate. Doubled haploid plants were obtained from haploid mid-vein cultures [19]. These plants were allowed to seed, and the seeds were used for further studies. Burley 49 transformed plants are referred to as NIa.

### 2.2. Transgene expression

Transgenic seed screening was performed as described before [7]. To detect the transcript containing TEV NIa sequences, total leaf RNA was isolated using phenol:chloroform. Denaturing formaldehyde gels were blotted onto nylon membranes and hybridized with antisense 32P-labeled RNA probes. Such probes were obtained by an *in vitro* cell-free transcription system of plasmid pTL37/8595, containing the NIa gene insert, using SP6 RNA polymerase.

### 2.3. Nuclear run-off assays

Isolation of nuclei from non-transformed and transgenic plant tissue was performed according to [20], with a slight modification in the amount of starting tissue (we used 3 g of leaves). DNA plasmids containing either the NIa insert, or the resident gene ubiquitin were digested using HindIII restriction enzyme. Then, 5 µg of the restricted products were electrophoresed in an agarose gel and blotted onto nylon membranes. Labelled 32P-RNA transcripts for run-off reactions were added in equal amounts (estimated by specific counts per million CPM) and used to hybridize the membranes containing DNA plasmids.

### 2.4. Inoculation of transgenic plants and non-transformed plants

In three independent replicates, 10 transgenic plants from each line were inoculated when they reached approximately 30 cm tall. Plant leaves were lightly dusted with carborundum and a 1:3 dilution of virus inoculum was applied with a cotton swab (dilution in w/v of virus-infected plant tissue in deionized distilled sterile water). Plant viral symptoms were recorded daily after the positive control plants started to show symptoms (8-10 days after inoculation). Six recovered leaves from transgenic plants and six non-recovered leaves from WT plants were inoculated with viral inoculum. This procedure was performed twice and surveyed for symptoms.

For the inoculation of wild type plants using transgenic stem or leaf tissues of recovered plants, we used the same dilution as before in six replicates and the experiment was repeated twice.

### 2.5 *In situ* hybridization

*In situ* hybridization was performed as described [21] with modifications. For synthesizing sense and anti-sense 11-digoxigenin-UTP labelled probes, a pTC plasmid containing a 1290-bp NIa fragment was linearized with restriction enzymes PsT I, and 1 μg was used as a probe synthesis template. RNA probes were synthesized using DIG northern starter kit (Roche, Mannheim, Germany). Transgenic non-inoculated (NIa-N plants) or TEV-inoculated and in process of recovery (NIa-PR1-PR3 and R) were fixed as described by [22]. RNA probes were hydrolyzed following the method described by [21], and 3%–6% of each labelling reaction (100–400 ng of RNA) was mixed with 40 μL of 50% formamide, added to 200 μL of hybridization buffer, and used as a probe for a pair of slides. Overnight hybridization was performed as described by [22].

### 2.6. Northern hybridization

Total RNA was resolved on a denaturing 0.8% agarose gel. Gels were transferred to a HybondN+ membrane (Amersham, Piscataway, NJ, USA). Hybridizations were performed with viral-derived probes labelled with a dCTP^32^P.

### 2.7. RNA extraction and Illumina Sequencing

Total RNA extraction was performed as described by [23]. Briefly, tissue was collected from leaves or stems from 4 plants in the process of recovery of PR3 and grounded in liquid nitrogen. A total of 4 RNA-Seq libraries (2 tissues, 2 replicates per tissue 2x100) were prepared and sequenced at Labsergen facilities (Irapuato, Mexico) using Illumina HiSeq2500 (Hayward, CA, USA). Raw data is publicly available at NCBI, BioProject accession: PRJNA844420.

### 2.8. Mapping and statistical analysis

Trimming and cleaning of reads were conducted according to [24]. Cleaned and trimmed reads were aligned against the reference tobacco transcriptome Nitab-v4.5 (www.solgenomics.net) [25,26] to quantify the expression of all transcripts using Kallisto v v0.46.2. Kallisto was executed with bias correction and 1000 bootstrap. Differential expression analysis was conducted with Sleuth v0.30.0 [27]. Reads of each library were mapped with a percentage of 81.05% pseudoaligned and 80.75% pseudoaligned just once to the transcriptome (Table S1). After the differential expression analysis, transcripts with log2 FoldChange (log2FC) values of ≥1 and ≤-1 and q-value of ≤0.05 were considered as differentially expressed genes (DEGs).

### 2.9. Functional annotation and gene expression validation

To identify the Gene Ontology enrichment, Singular Enrichment Analysis (SEA) was performed with AgriGO v2 with the Nitab4.5 (www.solgenomics.net) background under Fisher and the multi-test adjustment method of Hochberg (FDR) with a p-value ≤0.05 [28]. Ontology reduction and network construction were performed with REVIGO with standard parameters (medium 0.7 resulting list) and Cytoscape v3.8.0 [29,30]. Validation of gene expression and the presence of CP (viral) and NIa (viral and/or transgene) RNAs in leaves or stems was performed using quantitative reverse transcription PCR (qRT-PCR). The experiments were conducted on a StepOne Real-Time PCR System (Applied Biosystems) with qPCR SybrMaster HighROX (Jena Bioscience, Germany). The qRT-PCR was carried out under a three-step PCR protocol, with an alignment step at 60°C for 30 seconds.

The relative quantification of gene expression was determined using the double delta Ct (2^-ΔΔCt^) method, with actin serving as the housekeeping gene for normalization. The specific primers used for amplifying the different genes are listed in Table S2.

### 2.10. Small RNA analysis

A total of 4 small RNA-Seq libraries (2 treatments, 2 replicates per treatment 1x36) were prepared with gel band selection and sequenced at Cinvestav facilities (Irapuato, Mexico) using the Illumina HiSeq2500 (Hayward, CA, USA) platform. Raw data are publicly available at NCBI, BioProject accession: PRJNA844411. Reads were mapped with Bowtie v1.2.3 (--best --strata parameters) to the tobacco and TEV concatenated genomes, then miRNA prediction was carried out with ShortStack v3.8.5 standard parameters only quantifying miRNAs annotated in the miRBase database [31]. Differential expression analysis was carried out with DeSeq2 v1.36.0 standard parameters, and the target prediction of differentially expressed miRNA was carried out using psRNATarget 2017 V2 schema standard parameters [32].

### 2.11. Callose deposition analyzed by aniline blue multiphoton microscopy

Sections of 3 leaves and stems of NIa-plants at the PR3 and mock3 stage, and WT I3 and mock3 were cut and fixed in 4% formaldehyde, faded with 96% alcohol, and processed after the addition of SynaptoRed C2 as stated by the manufacturer (Biotium, Fremont, CA, USA), aniline blue staining was carried out following the methodology described by [33]. Multiphoton images were taken with an LSM 880-NLO Axio Imager Z2 microscope (Zeiss, Germany) coupled to a Ti:Sapphire infrared laser (Chameleon Vision II, COHERENT, Scotland). The laser wavelength was set at 723 nm for aniline blue and 800 nm for SynaptoRedC2, both dyes were detected at a 300-685 nm wavelength window with a Non-Descanned Detection.2 (NDD.2) using a 500-550 nm filter (Zeiss, Germany). Callose quantification was performed as described by [33].

## 3. Results

### 3.1. NIa-transgene expression and recovery

Following TEV infection, transgenic tobacco plants expressing either an untranslatable or a translatable version of the TEV NIa cistron (Figure S1) developed a systemic infection characterized by mosaic symptoms in the inoculated leaf and the three leaves immediately above the inoculated one (Figure 1). The process of recovery (PR) assessed as a symptom reduction was first observed in the interveinal regions of the leaf above the inoculated one (Nla-PR1) as well-defined chlorotic areas or speckles in interveinal regions. Between 14 to 21 days post-inoculation (dpi), the PR progressively increased in leaves 2 (NIa-PR2) and 3 (NIa-PR3), *i.e.,* NIa-PR1< NIa-PR2< NIa-PR3, registering an almost complete recovered leaf 4 (NIa-R4) (Figure 1a) sometimes with a reduced chlorotic speckle at the tip of the leaf. Once recovered, the leaves remained highly resistant to the virus as they showed no viral symptoms compared to WT plants. In contrast, all the leaves of WT plants remained symptomatic, showing chlorosis, while those of mock-inoculated NIa tobacco (NIa-N1-4) developed normally (Figure 1b and c, respectively). As expected, no infection was observed when fully recovered leaves were used as a source of inoculum in healthy plants (Figure S2).

**Figure 1:**
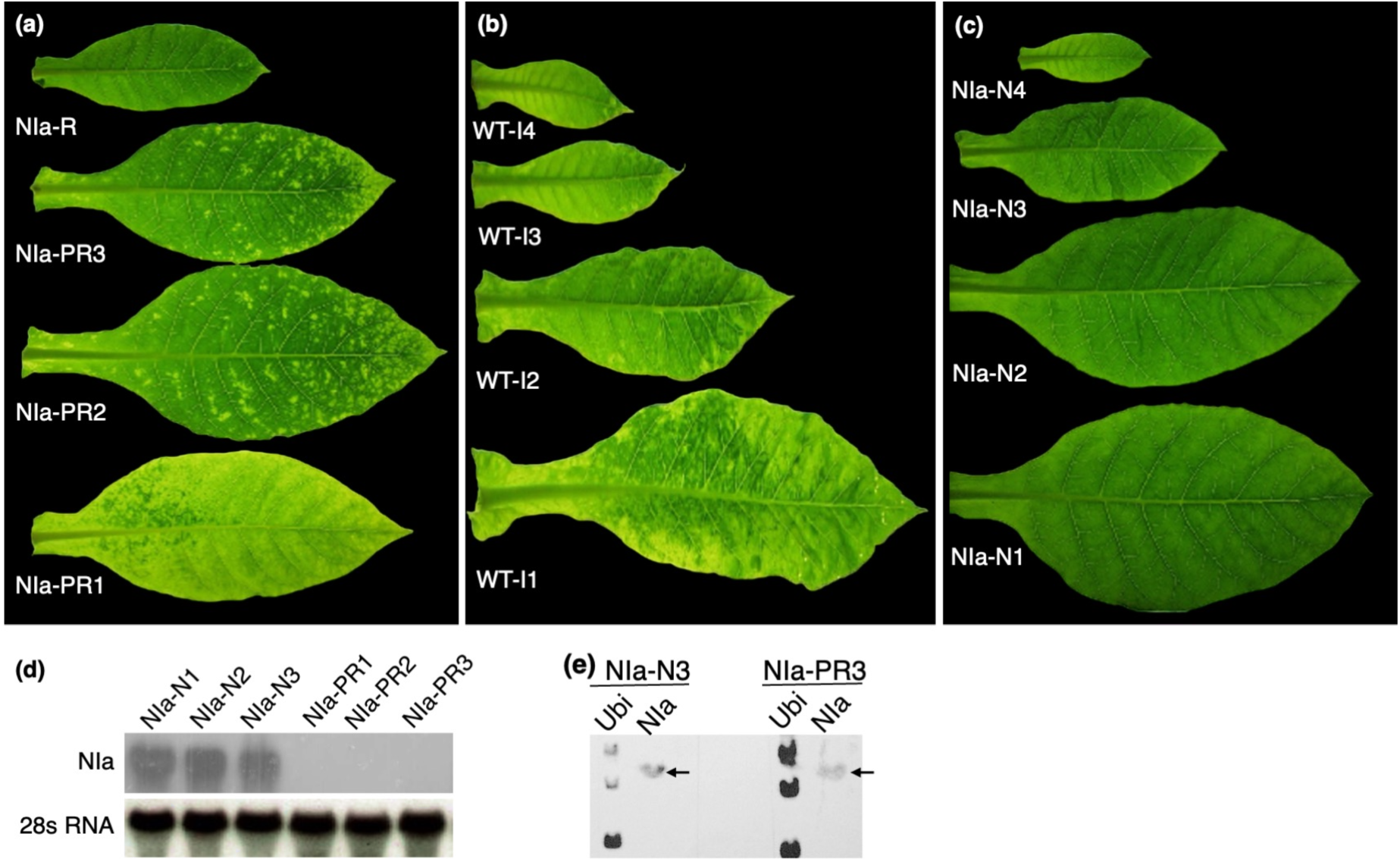
RNA silencing is involved in progressively recovering leaves from NIa plants. **(a)** Inoculated NIa plants in the process of recovery (NIa-PR1-3) and fully recovered (NIa-R) **(b)** Inoculated B49 wild-type plants (WT-I1-4). **(c)** The representative phenotype of mock-inoculated NIa plants (NIa-N1-4). **(d)** Northern blot of the NIa viral transgene-derived transcripts and 28S RNA (mostly cytoplasmic) of NIa-N1-3 and NIa-PR1-3 leaf tissue. **(e)** Run-off assay of the endogenous plant ubiquitin and viral transgene-derived transcripts in the extracted nucleus in leaves of NIa-PR3 and NIa-N3.

### 3.2 PTGS is involved in transgenic NIa-tobacco recovery

We confirmed that the PR observed in the leaves of NIa plants resulted from PTGS. By Northern blot, the expected band corresponding to the NIa RNA transcript is detected in NIa-N1-3 but not in NIa-PR1-3 leaves (Figure 1d). However, nuclear transcription rates of the transgene-derived NIa cistron and the endogenous ubiquitin gene were similar in mock-inoculated and recovered tissue at a similar physiological stage (NIa-N3 and NIa-PR3 at 21 dpi, integrated density values, IDV = 1.99 and 2.20, respectively). The multiple bands in the ubiquitin control lane resulted from incomplete digestion of the plasmid blotted onto the membrane (Figure 1e). This indicated that NIa transgene expression is similar in both the nuclei of plants in the process of recovery and mock-inoculated plants, and that recovery and viral gene silencing were activated following virus replication in the cytoplasm. The presence of the viral transgene-derived transcript in the nucleus and its absence in the cytoplasm suggested that PTGS was active during the recovery process.

### 3.3 Stem and leaf-vascular tissue did not undergo silencing as shown by *in situ* hybridization

We used the antisense viral NIa segment as an RNA probe to detect viral and/or transgene-derived transcripts in the leaf blade and leaf stem base, *i.e.,* the area of the stem that connects to the petiole, of NIa-N and NIa-PR plants by *in situ* hybridization (Figure 2). A positive signal in NIa-N tissues indicated the presence of the transgene-derived transcript. In contrast, the corresponding signal in NIa-PR tissues indicated the presence of viral transcript due to infection or the transgene-derived transcript. In the leaf blade, we observed transgene-derived transcript in NIa-N1-3 leaves (Figure 2a), whereas very faint or no signal was detected in NIa-PR leaves (Figure c). This observation indicated RNA silencing of both the viral-derived transcript and the virus. Conversely, viral and/or transgene expression at the stem was observed in both mock and viral inoculated NIa plants (Figure 2b and d). In mock plants, we observed a strong transcript signal in all cell types, including upper and lower epidermis, palisade, and spongy parenchyma, with higher intensity in the nucleus. Vascular bundles also presented a strong transcript signal. In the stem, phloem cells (companion cells and sieve elements) showed evidence of the transgene-derived transcript (Figure 2a-b). In some areas, phloem parenchyma cells were also indicative of the presence of the transcript. The brown color is due to non-specific background signal.

**Figure 2:**
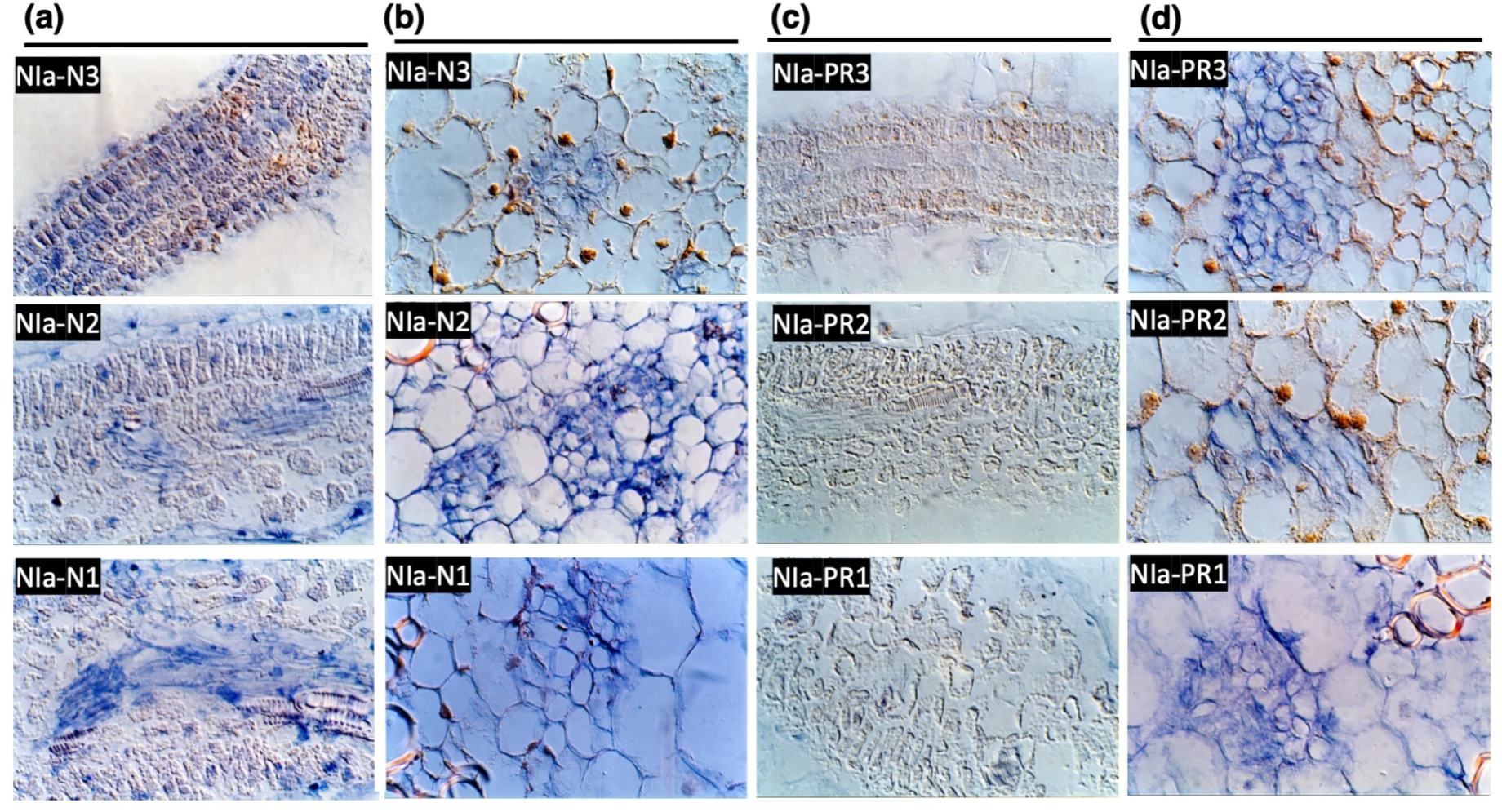
RNA silencing occurred in leaves but not in stems of infected NIa plants. The presence or absence of TEV-NIa RNA derived from the transgene and/or viral transcript was analyzed by *in situ* hybridization. **(a)** Leaves and **(b)** stems of transgenic mock-infected plants (NIa-N1-3). **(c)** Leaves in the process of recovery of transgenic infected plants (NIa-PR1-3) and **(d)** the corresponding sections of their stems.

Stem and leaves cut sections of NIa-PR3 were used as the source of inoculum in healthy WT plants; those inoculated with NIa-PR3 leaves remained symptomless. Meanwhile, plants inoculated with NIa-PR3 stems developed a symptomatic TEV infection (Figure S2). Furthermore, qRT-PCR carried out on CP RNA (viral) and NIa (viral and/or transgene) did not amplify the CP on NIa-PR3 leaves (Figure S3). These results validate the *in-situ* hybridization observation of absence of TEV in leaves and its presence and/or the NIa derived transcript in stems.

As expected, mock-inoculated WT B49 plants showed no evidence of viral expression in the leaf or stem. In contrast, the inoculated counterparts showed hybridization profiles like mock-inoculated NIa plants, albeit with a slight reduction of virus signal in leaves 2 and 3 (Figure S4).

The boundary zone at the stem-petiole insertion presented a differential accumulation of signal derived from the viral transcript of NIa (Figure 3). For instance, NIa RNA accumulated in the stem rather than in the petiole, except for the petiole vasculature, where NIa RNA was present (Figure 3a). Likewise, NIa was present in the leaf vasculature but not in the mesophyll tissue when we analyzed longitudinal sections of the leaf (Figure 3b).

**Figure 3:**
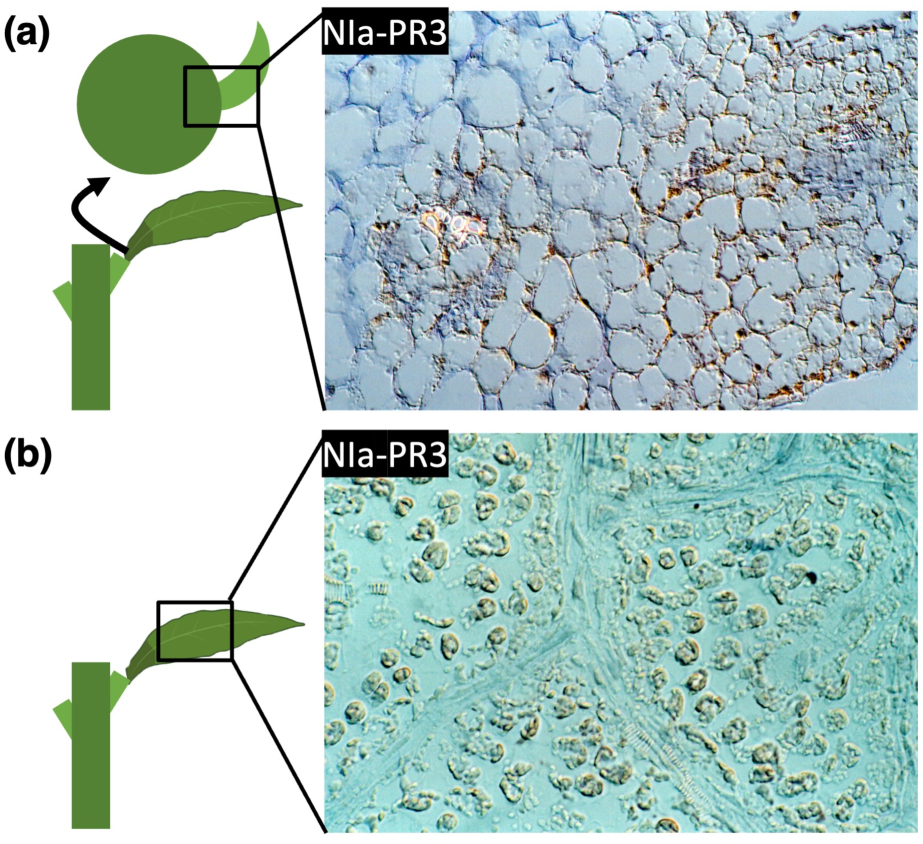
Vicinity between silenced leaf with non-silenced vasculature and non-silenced stem. *In situ* hybridization of RNA of tobacco TEV-NIa plants at the: **(a)** junction of stem-petiole insertion of NIa-PR3 plant and **(b)** leaf section of an NIa-PR3 plant showing non-silenced vasculature and silenced mesophyll cells.

### 3.4. Defense-related genes were overexpressed in leaf

To explore leaf and stem-specific responses to TEV during recovery, we performed high throughput RNA sequencing using transgenic NIa-plants at the PR3 stage. We identified 731 differentially expressed genes (DEGs) (log2FC ≥1 and ≤-1 and q-value of ≤0.05); among them, 700 DEGs corresponded to the leaf and 31 DEGs to the stem. The identified DEGs were enriched for functions related to the “binding” molecular function (MF) in the stem. On the other hand, 16 enriched biological processes (BP) were identified in the leaf, including “defense response”, “regulation of RNA metabolism”, “protein phosphorylation”, and “ubiquitination” (Fig. S5). Similarly, 17 MFs including “oxidoreductase activity”, “protein binding”, “sequence-specific DNA binding”, and “transcription factor activity” were enriched in this tissue. Overall, we found 28 disease-resistance, 8 jasmonic acid, and 3 ethylene-related genes differentially expressed in the leaf (Table S3), whereas no defense-related genes were differentially expressed in the stem.

### 3.5. The silencing machinery was overexpressed in the leaf, while suppressors of silencing were overexpressed in the stem

As silencing has been reported as the principal mechanism of recovery, we analyzed nine key silencing genes (DCL2, DCL4, SGS3, HEN1, NRPD1a, RDR6, RDR2, AGO1, and AGO2) differentially expressed. We found that four of the nine key RNA silencing genes, namely AGO1, DCL2, HEN1, and RDR6, were overexpressed in the leaf compared to the stem (Table 1). The other five genes were not differentially expressed. Moreover, in the stem, we found one transcript encoding for the calmodulin-binding protein-like (rgs-CML), a virus-triggered endogenous suppressor of RNA silencing well-characterized in tobacco (Table 1).

**Table 1.**
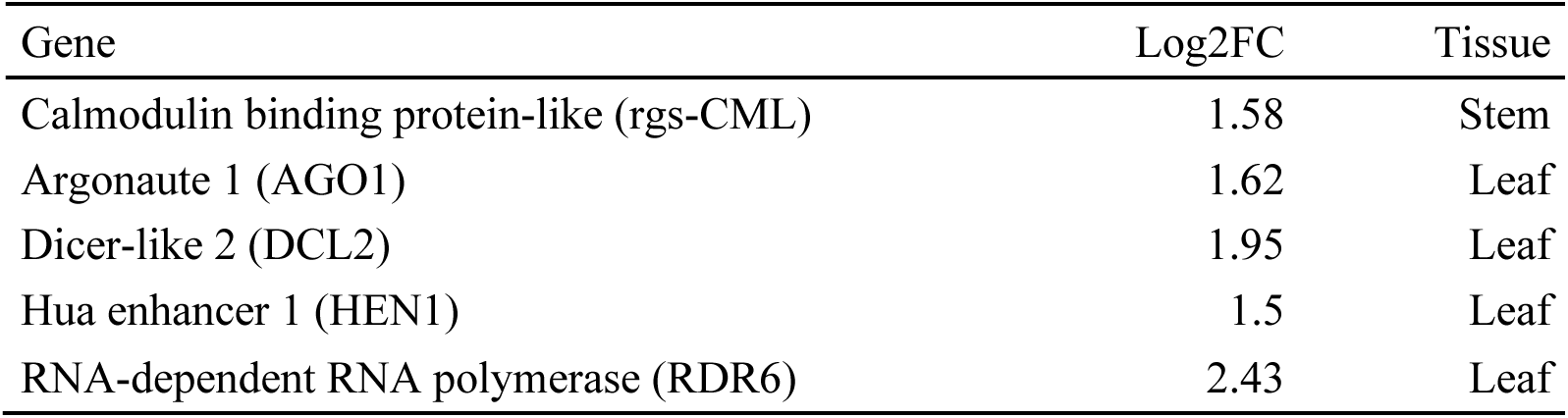
Silencing and repressor of gene silencing related DEGs expressed at leaf or stem.

| Gene | Log2FC | Tissue |
| --- | --- | --- |
| Calmodulin binding protein-like (rgs-CML) | 1.58 | Stem |
| Argonaute 1 (AGO1) | 1.62 | Leaf |
| Dicer-like 2 (DCL2) | 1.95 | Leaf |
| Hua enhancer 1 (HEN1) | 1.5 | Leaf |
| RNA-dependent RNA polymerase (RDR6) | 2.43 | Leaf |

### 3.6. Callose accumulation in the leaf suggested restriction of viral cell-to-cell movement by plasmodesmata

We further analyzed differences in the expression patterns of genes involved in viral movement across tissues. Notably, for the leaves we found we found overexpression of genes encoding for enzymes related to callose synthesis (three glucan synthase-like enzymes -GSL, and one callose synthase -CALS), and five genes related to callose accumulation at plasmodesmata (Table 2). The expression levels of rgs-CML, DCL2, AGO1, RDR6, GSL12 and CALS1 were validated by qRT-PCR (Figure S6).

**Table 2.** Callose synthase and deposition at plasmodesmata DEGs expressed in the leaf.

| Gene | Log2FC |
| --- | --- |
| GLUCAN SYNTHASE-LIKE 12 (GSL12) | 4.58 |
| CALLOSE SYNTHASE 1 (CALS1) | 2.85 |
| CADMIUM SENSITIVE 1 (CAD1) | 2.08 |
| GLUCAN SYNTHASE-LIKE 1 (GSL1) | 1.99 |
| IRREGULAR XYLEM 1 (IRX1) | 1.43 |
| PENETRATION 3 (PEN3) | 1.42 |
| GLUCAN SYNTHASE-LIKE 3 (GSL03) | 1.2 |
| UDP-GLUCOSYL TRANSFERASE 74B1 (UGT74B1) | 1.17 |
| CELLULOSE SYNTHASE A4 (CESA4) | 1.11 |

Following these results, we decided to stain NIa-PR3, NIa-mock3, WT-I3 and WT-mock3 leaves and stems with aniline blue to corroborate callose deposition at NIa-PR3 leaf plasmodesmata. In NIa-PR3 leaves, we found a significantly enhanced callose deposition compared to NIa-mock3, and callose accumulation was higher in NIa-PR3 leaves than WT mock or infected leaves (Figure 4a-b). In NIa-PR3 stems, callose accumulation was reduced in comparison with the other samples. However, this difference was not significant. We found a significant reduction of callose accumulation in infected leaves of WT plants, but no difference was shown between the infected and mock stems (Figure 4). These results corroborate the overexpression of callose accumulation-related genes at NIa-PR3 leaves.

**Figure 4:**
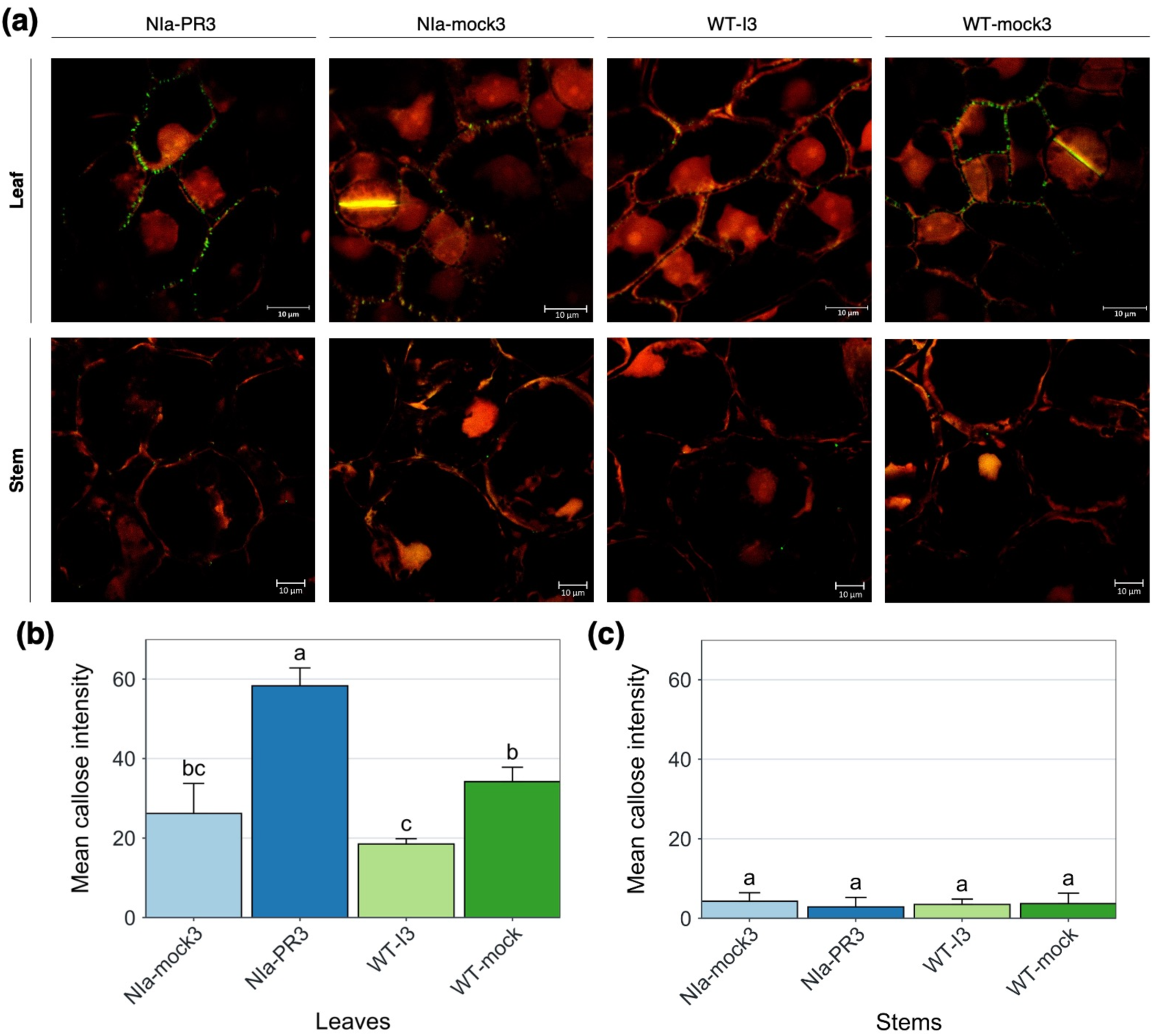
Plasmodesmata callose deposition in leaves and stems of NIa and WT plants. **(a)** Differences in callose deposition were observed using multiphoton imaging of aniline blue fluorescence. The red stain in the images corresponds to SynaptoRedC2 (plasma membrane staining), and the green color corresponds to aniline blue dye (callose staining). (b) Quantification of leaves callose deposits at plasmodesmata as observed after microscopy. (c) Quantification of stems callose deposits at plasmodesmata as observed after microscopy. Letters show grouping p-value <= 0.05 Tukey’s test.

### 3.7. Plant miRNA and vsiRNA profiles

A total of 12 miRNA were detected to be differentially expressed (p-value ≤0.05) in the leaf and 4 in the stem, with 77 and 6 predicted targets, respectively. Functional enrichment analysis of leaf miRNAs targets showed a negative regulation of macromolecule modification, protein phosphorylation, response to stimulus, response to stress, defense response and gene silencing by RNA (Fig. 5a) with miRNA targets of plant-pathogen interaction pathway and disease resistance (Table S4). Meanwhile, enrichment for stem miRNA did not show any functional enrichment.

**Figure 5:**
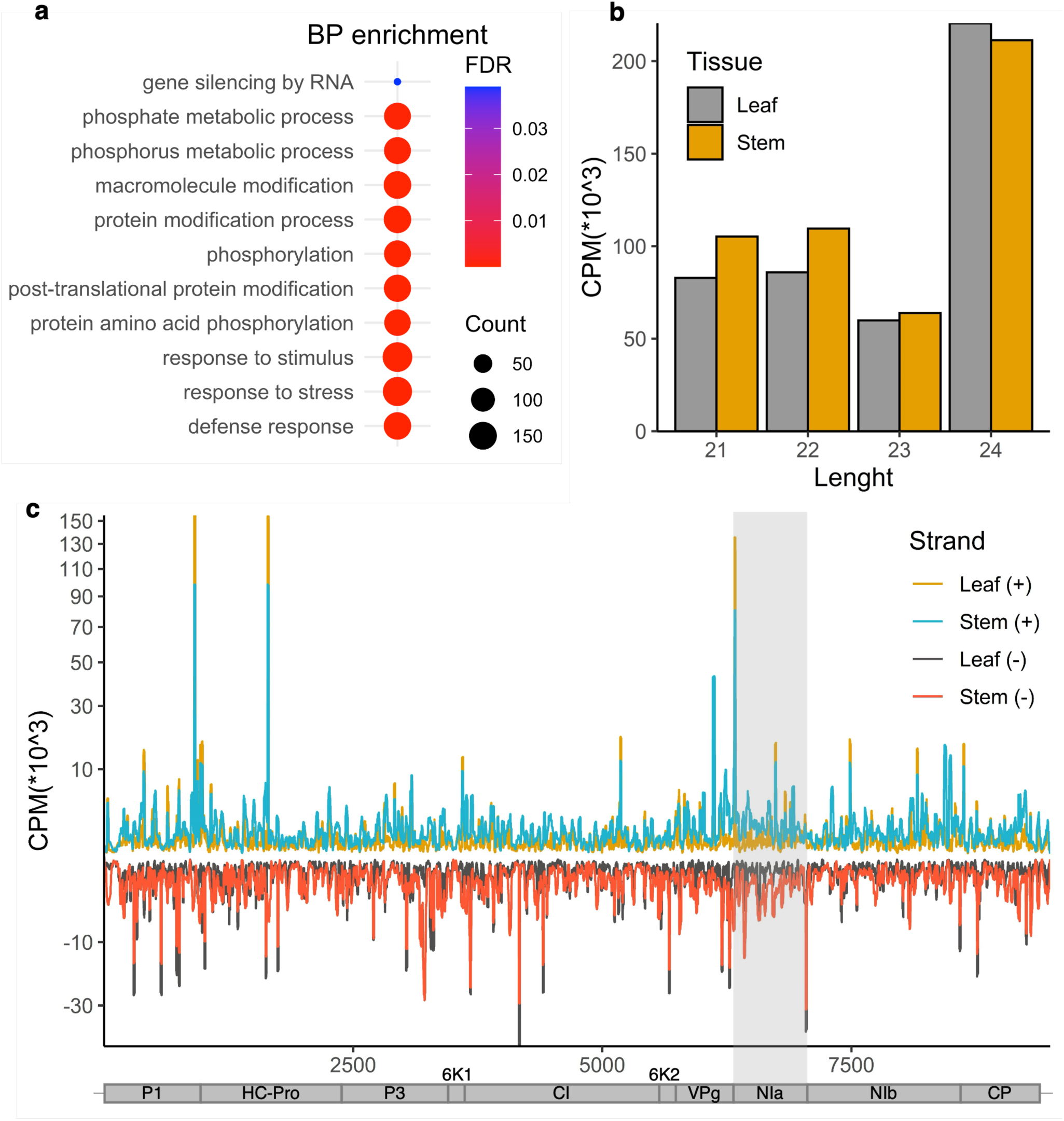
Plant miRNA and vsiRNA profiles in leaves and stem of NIa-PR3 plants. **(a)** Functional enrichment network of Leaf miRNAs. **(b)** Per tissue abundance of 21, 22, 23, and 24 nt vsiRNA. **(c)** TEV genome-wide distribution of vsiRNA in counts per million (CPM).

Virus-derived small interfering RNAs (vsiRNAs) are commonly produced during viral infections and work as keystone molecules for effective viral silencing. In this study, we observed that vsiRNA was more abundant in the stem than in leaves (489.89 CPMx10^-3^ and 448.96 CPMx10^-3^, respectively). However, 24 nt vsiRNAs were more abundant in the leaf, which could suggest a role for DCL3 in their biogenesis. Surprisingly, we did not observe differential expression of DCL3 in the leaf (Fig. 5b, Table 1). TEV genome-wide accumulation of vsiRNAs revealed *per-site* differences, with higher abundance at the P1 cistron than other genomic regions, including the NIa transgene (Fig. 5c). Log2Fold change was calculated to compare vsiRNA genome accumulation between leaf and stem. We found an overall lower vsiRNA accumulation in the leaf (Figure S7). In addition, we found remarkable differences in the negative-stranded vsiRNA of P3 and CI cistrons between leaf and stem, suggesting these RNAs could be important in maintaining a non-silenced stem (Figure S7b).

It has been proposed that vsiRNA has the potential to regulate host gene expression [34] Therefore, we investigated if vsiRNA derived from the top 20 abundant sites and P3 and CI cistrons target plant host genes. The resulting predicted target genes in the tobacco of both abundant and non-abundant vsiRNAs were not functionally enriched. However, we found that the stem-derived-vsiRNA arising from the P3 cistron region between ∼3200-3300nt (-) targets GSL03, a tobacco gene involved in viral movement restriction. Also, the most abundant vsiRNAs derived from the P1 and HC-Pro (∼900nt and ∼1600nt) region are predicted to target both DCL4 and PR1. No defense-related target genes were found in the non-abundant regions.

## 4. Discussion

### 4.1. Plants undergo a process of recovery which is compartmentalized

We suggest that the process of recovery in leaves of NIa plants resulted from PTGS, as in other recovery systems [3]. Recovered tissues remain resistant to the virus, as reported before in other transgenic tobacco plants expressing the TEV-viral coat protein cistron [17]. The presence of the virus in the stem and leaf vasculature but not in the mesophilic cells suggested restriction of intracellular movement of TEV, possibly occurring at the exit from the phloem. Previous studies have shown that not all the new emerging organs are fully recovered, as we saw between the recovered leaves and non-recovered stems in the present study. However, none of these works compared the differences between recovered and non-recovered tissues [35,36].

### 4.2. RNA silencing and viral movement restriction at leaf and stem

It is well known that RNA silencing plays a key role in recovery, as the elimination of components of the RNA silencing machinery halts plant recovery [35,37]. Viruses that efficiently counteract RNA silencing evade recovery [38,39]. Here we found that genes that are essential for RNA silencing, including AGO1, DCL2, HEN1, and RDR6, were overexpressed in the leaf but not in the stem. Interestingly, AGO2 was not differentially expressed, albeit previously described as the main silencing gene involved in antiviral defense [12]. Similarly, other key genes for VIGS and siRNA biogenesis, such as SGS3 and SDE5, and key genes related to PTGS-TGS crosstalk, such as NRPD and RDR2, were not differentially expressed in the stem [37,40,41]. These findings suggest that other factors may be involved in the stem repression of silencing since some key silencing-related genes are equally expressed in both tissues (DCL3-4, SGS3, NRPD1a, RDR2, and AGO2). In this regard, we found that the plant endogenous rgsCaM was overexpressed in the stem, suggesting it may participate in silencing suppression, as it is known to be recruited by TEV HC-Pro to suppress RNA silencing [42] and may explain the lower expression of RDR6 [43]. Moreover, rgsCaM is also known to enhance virus resistance by targeting itself and HC-Pro to proteolytic pathways, thus restricting viral infections to local areas [44]. In this sense, the overexpression of rgsCaM could explain the lack of silencing in the stem and enhancement of antiviral responses restricting viral infection to this tissue.

Another point to consider is that RNAi components could be transported from the leaves to the stem based on local RNA expression levels. While RNA silencing genes were not overexpressed in the stem, this does not necessarily mean their proteins or RNAi products are inactive in this tissue. It is plausible that RNAi elements, such as siRNAs or silencing-associated proteins, are shuttled from the leaves to the stem, contributing to antiviral defense. Further studies are needed to determine whether RNAi components generated in the leaves are transported to the stem and how they function in regulating viral silencing and movement across tissues [45–47].

### 4.3. Plant innate immunity also participates in plant recovery

The lack of tissue recovery in tissues such as the meristem has been attributed to RNA silencing and is considered a key step in viral recovery [48]. Recent works provide evidence that silencing is not the reason for tissue exclusion of recovery, as DCL2/4-silenced plants infected with CymRSV were free of the virus at the meristem [49]. Most of the literature describes recovery by specific components in the RNA silencing pathway rather than considering the global changes occurring in a plant during the recovery process. PAMP-triggered immunity (PTI) has been reported to be prevalent in almost all plant-virus transcriptomes. Indeed, we detected many genes involved in PTI and canonical defense overexpressed in the leaf. For instance, nine genes related to callose deposition at plasmodesmata or negative regulators of plasmodesmata permeability (GSL12, GSL1, GSL03, CALS1, CAD1, IRX1, PEN3, UGT74B1, and CESA4) were overexpressed in the leaf. In agreement with these observations, we corroborated callose accumulation by image microscopy in the leaf. Similar to our findings, a Cassava line resistant to CBSV allowed viral transmission through the stem, whereas no virus could be detected in leaves [50]. In the following study, the authors found overexpression of b-1,3-GLUCANASE, ANKs, and PDLP1 genes in the susceptible line. These genes participated in plasmodesmata permeability as the viral movement-promoting factors, whereas GSL4 (a possible negative regulator of plasmodesmata permeability) was overexpressed in both susceptible and resistant lines [51]. However, we did not find any viral movement promotor overexpressed in the stem, which suggests that, in our system, viral movement restriction is exerted through a pathway different to the one identified in CBSV-resistant Cassava.

### 4.4. vsiRNA accumulation at leaf and stem

Contrary to the findings in ORMV-infected Arabidopsis plants, which had a high amount of 21-22nt vsiRNA accumulated in recovered actively virus-infected leaves [37], in this study, we quantified around 20% less 21-23nt vsiRNAs in recovered tissue (leaf) than in non-recovered stem. Out of the 7 genes dubbed crucial for recovery [37,41] four (AGO1, DCL2, HEN1, and RDR6) were overexpressed in the leaf compared with the stem. The higher accumulation of vsiRNAs at the stem could be explained by an active replication of the virus in this tissue. Despite the overexpression of DCL2 (involved in 22nt production) in leaves, we found more accumulation of 24nt than 21-23nt vsiRNAs. Because it is more common that 21-22nt vsiRNAs accumulate in higher amounts than 23-24nt vsiRNAs during viral infections [52,53], further investigation needs to be performed to explain our differential findings. We observed a higher accumulation of positive-strand vsiRNAs, which has been previously found to be common in positive-stranded RNA viruses [52]. The non-random distribution of vsiRNAs along the TEV genome is also common and is associated with the fold-back structures of the genomic RNA and binding of the vsiRNAs to the viral HC-Pro [54–56]. vsiRNA accumulation hotspots have been previously associated with the modulation of pathogenesis in the host machinery by alphavirus-derived small RNAs, which influence infection outcomes in disease vector mosquitoes [34]. However, in our work no vsiRNAs arising from any of the top 20 accumulation hotspots had predicted target genes involved in defense or the pathogenesis response in the host, including those arising from differential hotspots between leaves and the stem. But vsiRNAs do target GSL03 gene involved in viral movement restriction

## 5. Conclusions

Here we show that additional host mechanisms, such as the overexpression of defense-related genes and callose accumulation at plasmodesmata, contribute to viral disease recovery, complementing RNA silencing. We also bring insights that compartmentalization of RNA (viral of transgene derived) or specialization of recovery differentially occurs between leaves and stems since RNA silencing and plant defense responses are mounted differently in such tissues. It is also possible that RNAi components produced in the leaves may be transported to the stem, where they still contribute to antiviral defense, highlighting the importance of both local and systemic RNA silencing activity. Thus, this study opens the window to investigate whether defense responses innate and adaptive to viruses or other pathogens are also differentially triggered in other plant organs. Understanding this compartmentalization would allow the designing of new strategies to control diseases.

## Data availability

The raw reads generated during the current study are publicly available at NCBI under PRJNA844420, PRJNA844411 accession numbers. Materials are available from the corresponding author upon reasonable request.

## Supporting information

Supplemental talbles

Supplemental images

## Acknowledgments

We thank Gabriela Chávez-Calvillo, Gustavo Rodríguez-Gómez, Domitila J. Rosales, Dalia Rodríguez-Ríos, José N. Santoyo-Villa and Luis J. Saucedo-Vargas and PlanTecc for their technical support. This research was supported by grants from Cinvestav (IP 1995) and CONACyT (4274-N9406). LSR and SLFV supervised the study. LSR, PVM, and AOL designed the experiments. PVM, AOL and CMFG performed the experiments. PVM analyzed the data and wrote the manuscript. LSR, SLFV, GL and PVM edited the manuscript. JPVC directed the in-situ hybridization work performed in 2000. All authors read and approved the final manuscript. This paper is part of the productivity of the Doctoral degree of P.V.M. at the Posgrado en Ciencias Biológicas, UNAM.

