## Supplemental images for "Callose deposition at plasmodesmata suggests a role in viral movement restriction in transgenic tobacco plants during recovery from *Tobacco etch virus* infection"

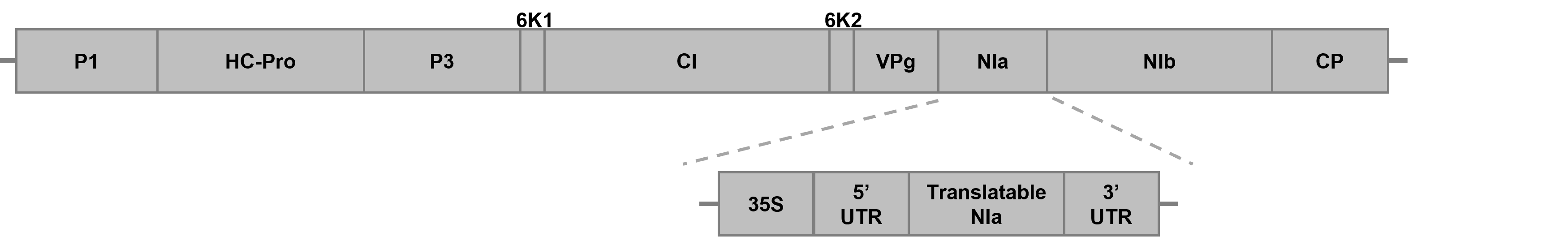

Figure S1: TEV NIa construct used for the transformation of tobacco plants.

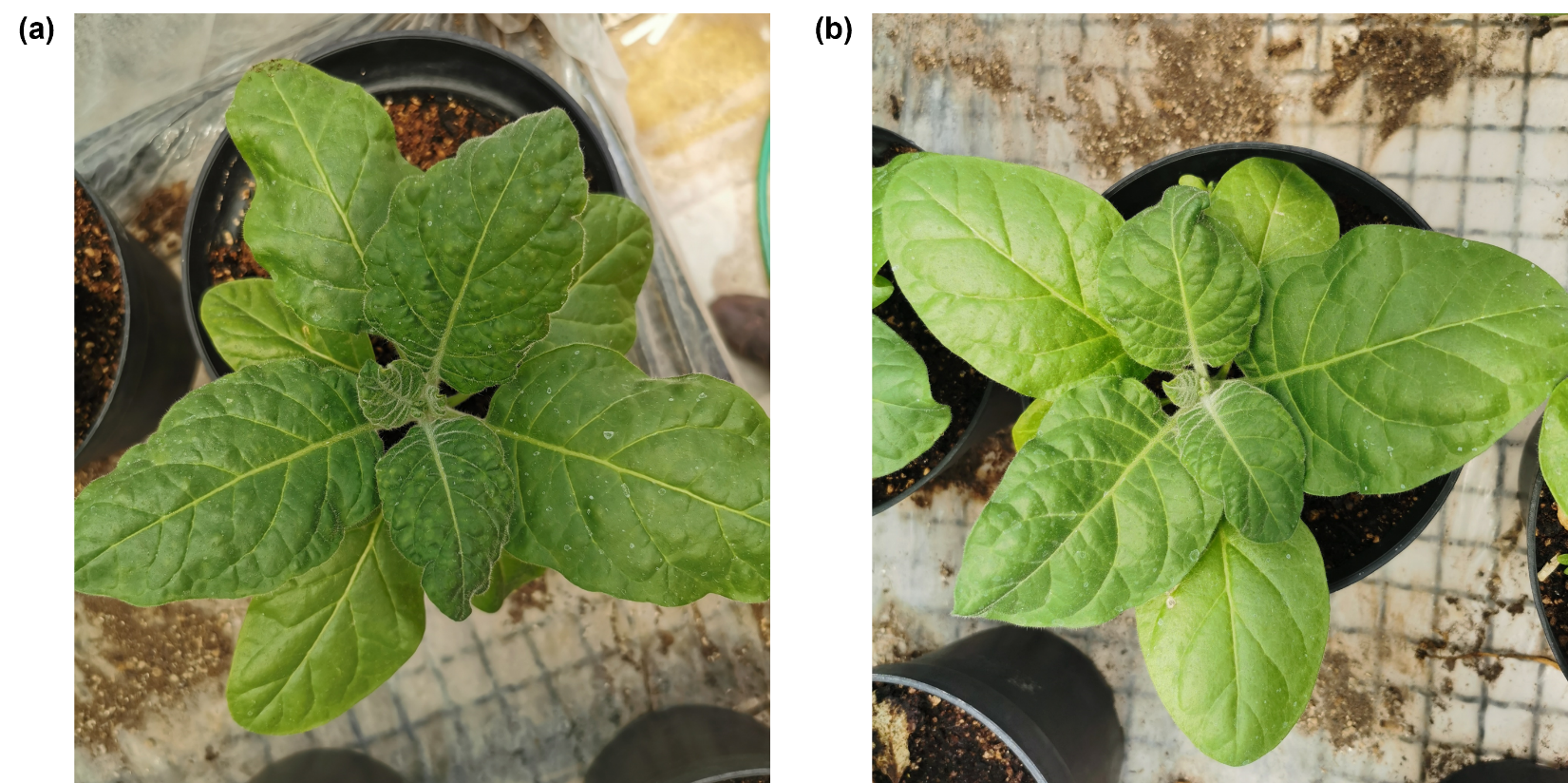

Figure S2: Stem remains infective but not recovered leaves. **(a)** Wild type tobacco plant infected with stems from a NIa-PR3 plant. **(b)** Wild type tobacco plant infected with a NIa-PR3 leaf.

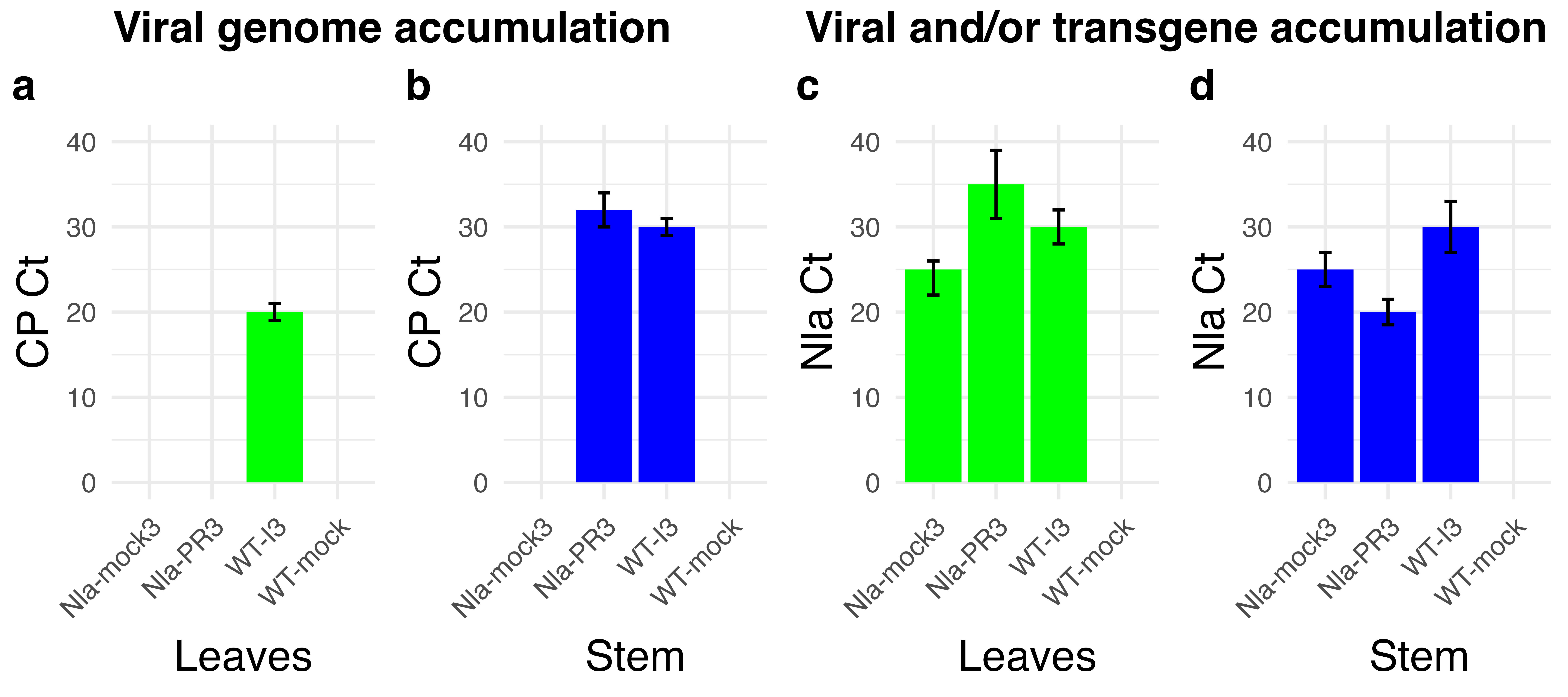

Figure S3: Presence of CP (viral) and NIa (viral and/or transgene in NIa viral in WT) RNA. Ct qPCR threshold cycle.

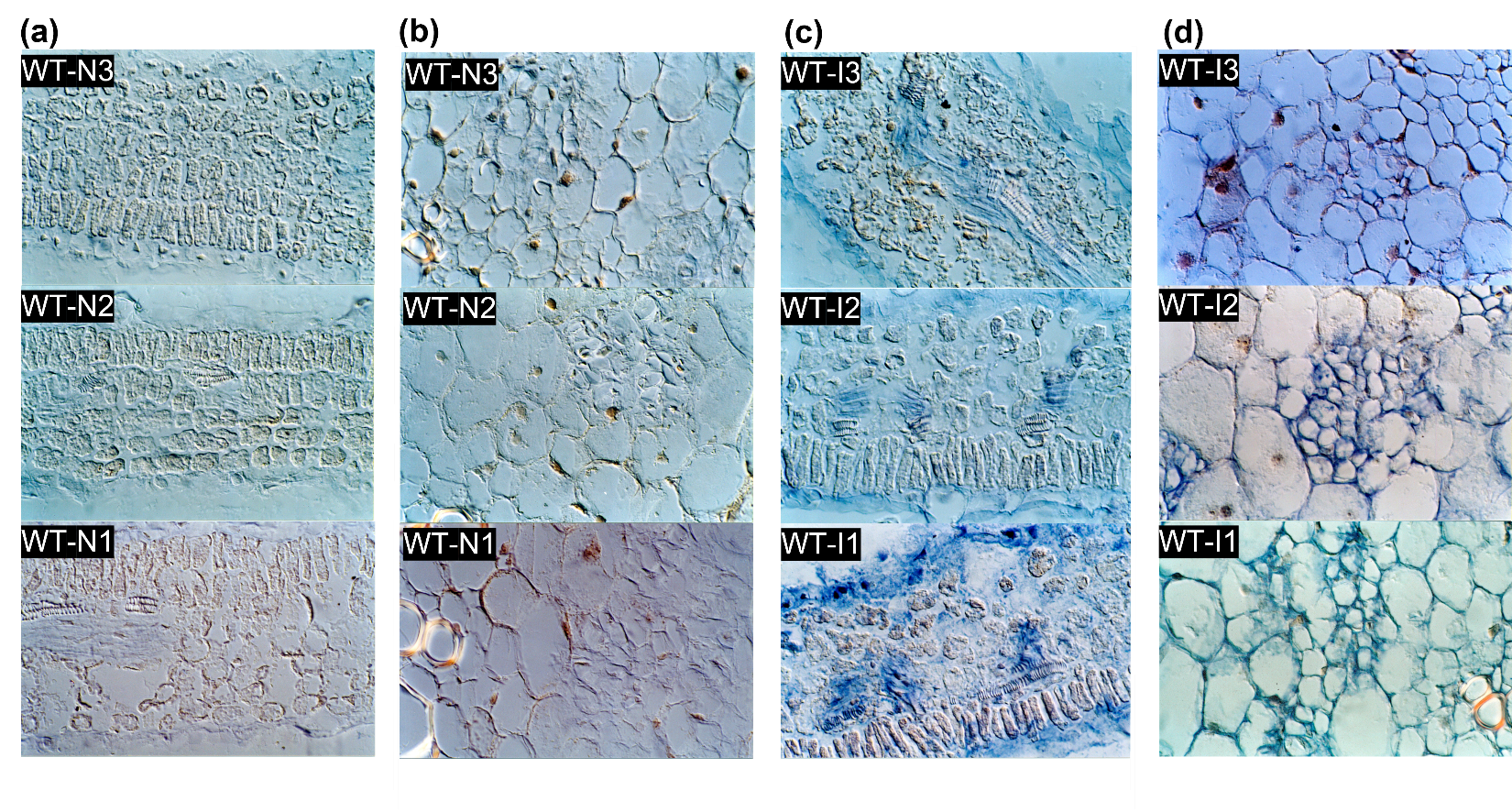

Figure S4: Presence or absence of TEV-NIa or viral derived transgene-transcript in uninfected and infected plants. Leaves **(a)** and stems **(b)** of wild type non infected tobacco plants (WT-N). **(c)** TEV infected wild type leaves (WT-I) and corresponding sections of insertion to the stem **(d)**.

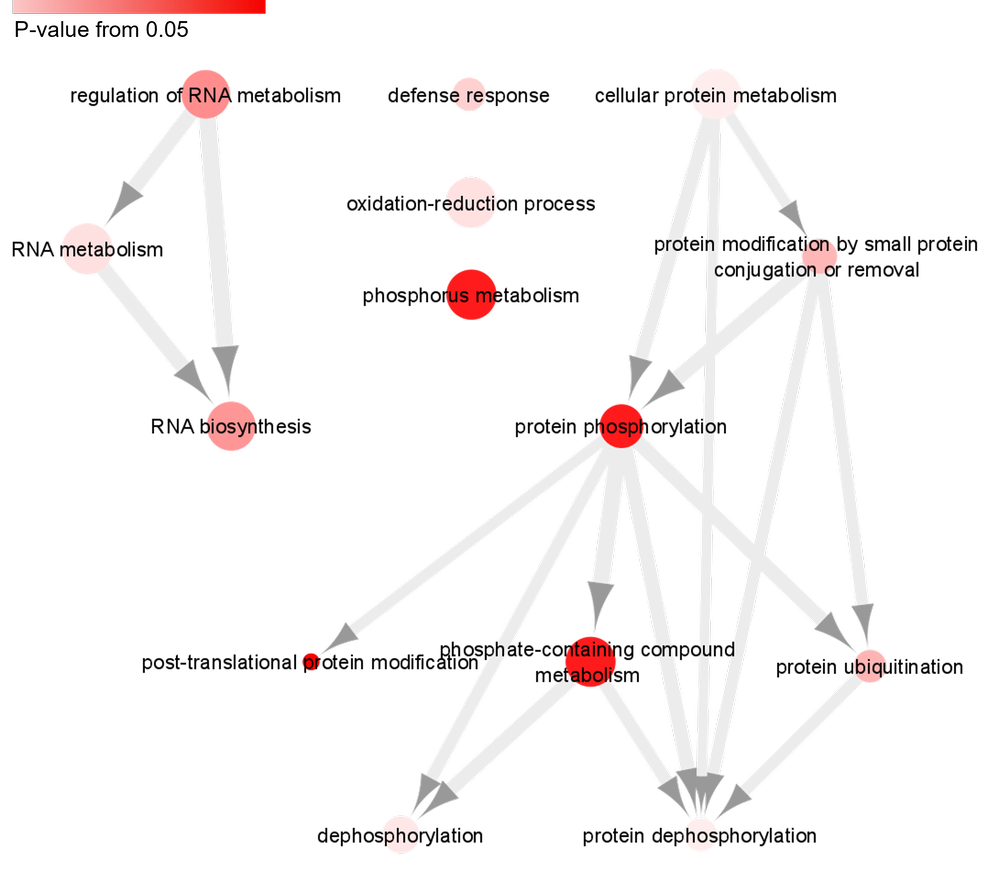

Figure S5: Upregulated biological processes in recovered leaves denoted through a functional DEGs enrichment analysis. Hierarchical network showing biological processes (BP) ontologies. Ontologies shown in the figure are only the semantically reduced terms in REVIGO. Edges represent relationships between the BPs (nodes) and edge weight represent the proportion of genes shared between enrichments. Intensity at node color indicates the p-value of the enrichment. The node size represents the number of genes at each enrichment.

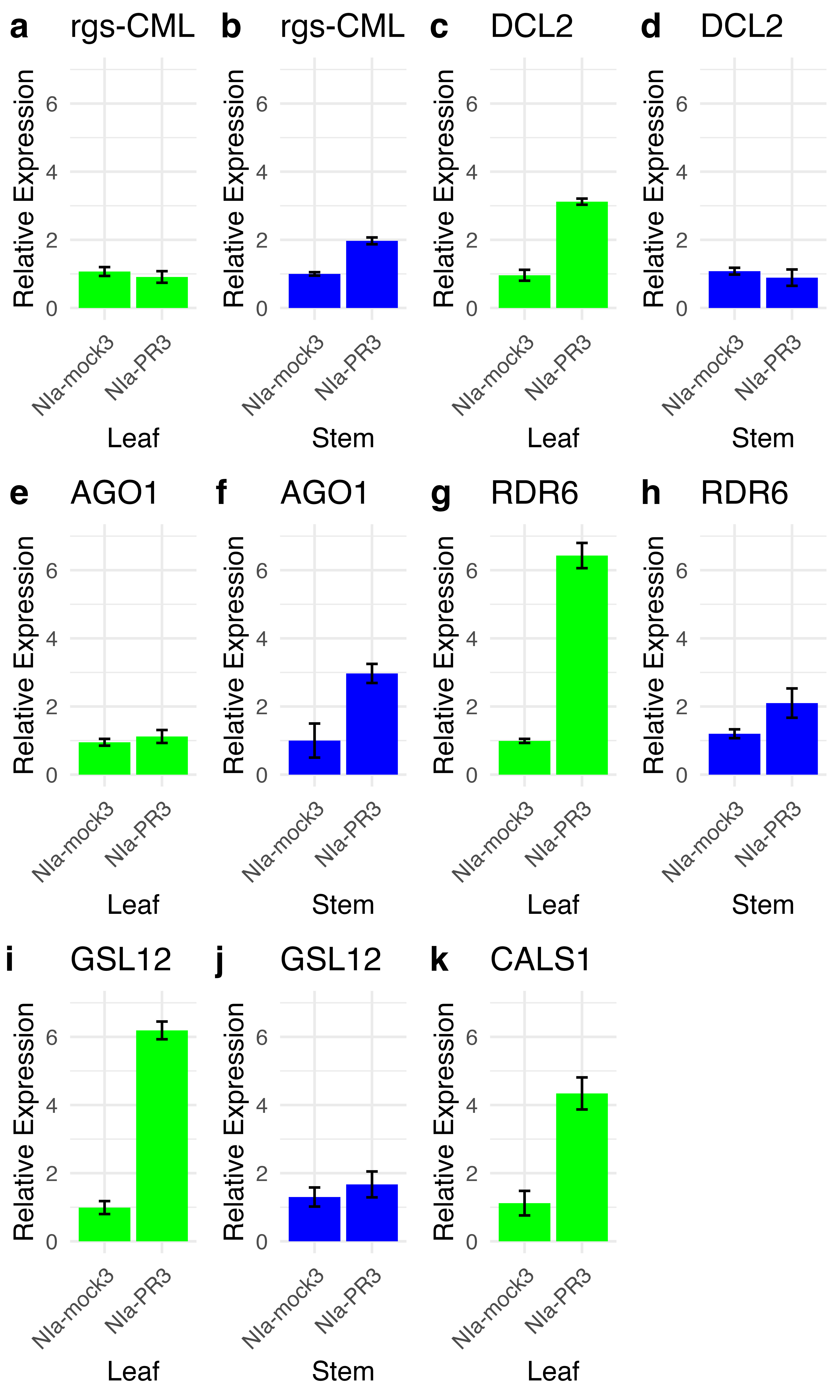

Figure S6: Relative expression of silencing and Callose synthase and deposition related genes. a-b Relative expression of rgs-CML in NIa mock and infected leaves and stems respectively. c-d Relative expression of DCL2 in NIa mock and infected leaves and stems respectively. e-f Relative expression of AGO1 in NIa mock and infected leaves and stems respectively. g-h Relative expression of RDR6 in NIa mock and infected leaves and stems respectively. i-j Relative expression of GSL12 in NIa mock and infected leaves and stems respectively. k Relative expression of CALS1 in NIa mock and infected leaves.

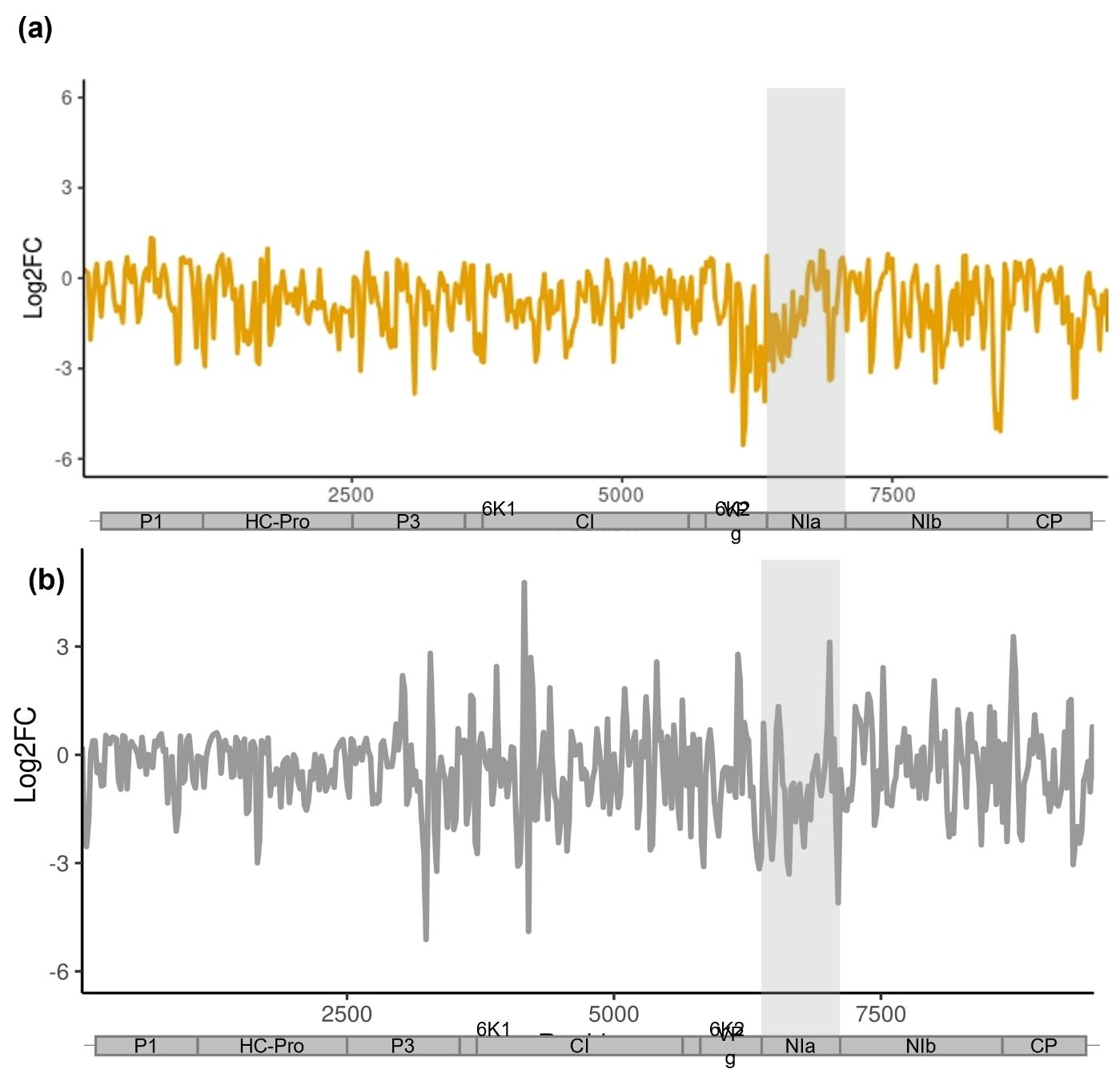

Figure S7: TEV genome-wide accumulation foldChange (Log 2 scaled) of vsiRNAs between leaf and stem. **(a)** Positive stranded vsiRNAs. **(b)** Negative stranded vsiRNAs.
